# PREVALENCE OF FUSARIUM WILT (*Fusarium oxysporum* f. sp. *Lycopersici*) on TOMATO (*Solanum lycopersicum* L.) IN CHIKUN LGA, KADUNA STATE

**DOI:** 10.64898/2026.06.24.734226

**Authors:** Olanrewaju Rilwan, Akos Ibrahim

## Abstract

Tomato (*Solanum lycopersicum* L.) is one of the most important vegetable crops in Nigeria, serving as a major source of income, nutrition, and raw material for food industries. However, its production is severely constrained by Fusarium wilt, a destructive soil-borne disease caused by *Fusarium oxysporum* f. sp. *lycopersici*. This study investigated the prevalence and severity of Fusarium wilt on tomato in Chikun Local Government Area (LGA) of Kaduna State, Nigeria. Field survey and laboratory analyses were conducted on forty-five tomato samples from three tomato farms Kujama, Kakau, and Rido. The samples were examined for disease incidence and severity. Data were analyzed using descriptive statistics and Chi-square tests. The overall disease incidence was with Rido recording the highest infection rate (80.0%), followed by Kujama (60.0%) and Kakau (40.0%). Among plant parts, the stem exhibited the highest infection frequency (80.0%), while leaves and fruits had 60.0% and 40.0% incidence respectively. Chi-square analysis indicated no significant difference (p > 0.05) in disease incidence among farms and plant parts, suggesting uniform pathogen distribution. The research recommends the adoption of integrated disease management strategies and improved farmer awareness to mitigate the impact of the disease and ensure sustainable tomato production.

## CHAPTER ONE

### 1.0 INTRODUCTION

#### 1.1 BACKGROUND OF THE STUDY

Tomato *(Solanum lycopersicum L.)* is one of the most important vegetable crops cultivated globally, valued for its nutritional, economic, and dietary contributions (FAO, 2022). In Nigeria, tomato production plays a significant role in food security and income generation, especially among smallholder farmers. However, the productivity of tomato has been significantly constrained by several biotic stresses, chief among them being *Fusarium* wilt caused by *Fusarium oxysporum f. sp. lycopersici* (Fol), a highly destructive soil-borne fungal pathogen (Abayomi et al., 2021).

*Fusarium* wilt is a vascular disease that impedes water and nutrient transport in tomato plants. This soil-borne fungal pathogen invades the plant through its roots, colonizes the vascular system, and ultimately disrupts water and nutrient transport, leading to wilting, leaf yellowing, stunted growth, and eventual plant death (Okungbowa & Shittu, 2019). The disease is widespread in tropical and subtropical regions, with increasing prevalence reported across various agro-ecological zones in Nigeria (Olorunjuwon et al., 2020).

Fusarium wilt is particularly problematic in tropical and subtropical regions such as Nigeria, where warm temperatures and poor soil management practices create conducive environments for pathogen persistence and spread (Agrios, 2005). The prevalence and severity of the disease have led to significant yield losses, estimated to range from 30% to 80%, depending on environmental conditions, cultivar susceptibility, and disease management strategies employed (Adebola & Afolayan, 2020).

The pathogen’s ability to survive in the soil for long periods, its variability in virulence, and its resistance to conventional control methods make it a persistent and challenging threat to tomato cultivation Haruna, *et al*., (2024). While chemical fungicides and resistant varieties have been explored, these approaches have shown limited success, especially under field conditions, and often raise concerns about cost-effectiveness and environmental impact (Abayomi, *et al*., (2021). Additionally, lack of reliable data on the current distribution and prevalence of FOL across tomato-growing regions in Nigeria’s diverse agro-ecological zones limits the development of sustainable, integrated disease management strategies (Olorunjuwon, *et al*., (2020). Moreover, the emergence of new races of the pathogen further complicates resistance breeding efforts and necessitates continual surveillance (Nwanguma et al., 2022). Understanding the current prevalence of *Fusarium* wilt in tomato-growing areas of Kaduna is critical to devising effective disease management strategies and supporting the development of resistant varieties. Although previous studies have documented the presence of the disease in some parts of the country, comprehensive and updated assessments remain limited. This study, therefore, seeks to determine the incidence and severity of *Fusarium oxysporum* f. sp. *lycopersici* in tomato fields in Chikun Local Government Areas in Kaduna, Nigeria. The findings will contribute to a better understanding of the epidemiological patterns of the disease and provide a scientific basis for integrated disease management practices.

#### 1.2 RESEARCH OBJECTIVES

i. To determine the incidence and severity of Fusarium Wilt (*Fusarium oxysporum* f. sp. *lycopersici*) on tomato (*Solanum lycopersicum* L.) in Chikun Local Government areas

#### 1.3 JUSTIFICATION OF THE STUDY

The increasing demand for tomatoes in Nigeria is challenged by declining yields due to *Fusarium* wilt, a persistent and destructive soil-borne disease that affects tomato roots and vascular systems. Smallholder farmers, who dominate tomato production, often lack access to reliable disease surveillance data and effective management options (Umar et al., 2020). The pathogen’s ability to remain viable in soil for several years and the lack of widespread use of resistant varieties contribute to its continued spread (Ezekiel et al., 2018). Additionally, climate variability and poor agricultural practices have further exacerbated the incidence of *Fusarium* wilt in tomato-growing regions (Ibrahim & Sani, 2021). A systematic study on the prevalence of the disease is therefore necessary to guide research, policy interventions, and extension services. Such data will also support the development of integrated disease management strategies and improve the resilience of tomato production systems in Nigeria.

#### 1.4 SCOPE OF THE STUDY

This study aims to determine the incidence and severity of *Fusarium* wilt, caused by *Fusarium oxysporum* f. sp. *lycopersici*, in Chikun Local Government Areas in Kaduna, Nigeria. The research involves field surveys to identify symptomatic tomato plants, sample collection, and laboratory-based pathogen confirmation using standard morphological techniques (Nwankiti, 2017). The study assesses disease incidence and severity across diverse agro-ecological zones, considering agronomic practices, soil types, and cultivar susceptibility (Okungbowa & Shittu, 2020). Although environmental and farming conditions contributing to disease spread will be examined, molecular characterization of pathogen races and long-term control efficacy are beyond the scope. This research is intended to provide baseline data necessary for the formulation of effective and location-specific management strategies, which are currently lacking in many parts of Kaduna. The outcome will support extension services and guide stakeholders toward improving tomato health and productivity.

## CHAPTER TWO

### 2.0 LITERATURE REVIEW

#### 2.1 Introduction

Tomato *(Solanum lycopersicum L.)* is one of the most consumed vegetables worldwide due to its nutritional, economic, and dietary value. In Nigeria, tomato farming plays a crucial role in food security and income generation, particularly in Kaduna State, where Chikun Local Government Area (LGA) is an important production zone. Despite its importance, tomato productivity is threatened by several diseases, with Fusarium wilt, caused by *Fusarium oxysporum f. sp. lycopersici* (Fol), being among the most devastating. This section reviews literature on the incidence and severity of Fusarium wilt (Agrios 2005).

#### 2.2 CHARACTERISTICS OF FUSARIUM WILT

Fusarium wilt of tomato is caused by *Fusarium oxysporum f. sp. lycopersici* (Fol), a soil-borne fungal pathogen that infects the vascular tissues of tomato plants. The disease is characterized by progressive wilting, yellowing of lower leaves, stunted growth, and eventual plant death (Agrios, 2005). One of its key features is the production of chlamydospores, thick-walled resting spores that enable the pathogen to survive in the soil for up to a decade even in the absence of host plants (Olorunjuwon, et al., 2020).

The fungus enters the plant through the roots and colonizes the xylem vessels, obstructing water and nutrient transport. This blockage causes vascular discoloration, visible as dark streaks in the stem when cut open, which is a diagnostic symptom of the disease Abayomi, *et al*., (2021). Fusarium wilt thrives in warm, moist conditions, with an optimum temperature of 25–30°C, conditions typical of Nigeria’s tomato-producing zones (Haruna, et al., 2024). The severity of infection varies with cultivar susceptibility, soil type, and cultural practices.

Overall, the persistence of the pathogen in soil, its ability to infect a wide range of tomato cultivars, and the absence of highly effective chemical controls make Fusarium wilt one of the most difficult tomato diseases to manage in Nigeria.

**Figure 1:**
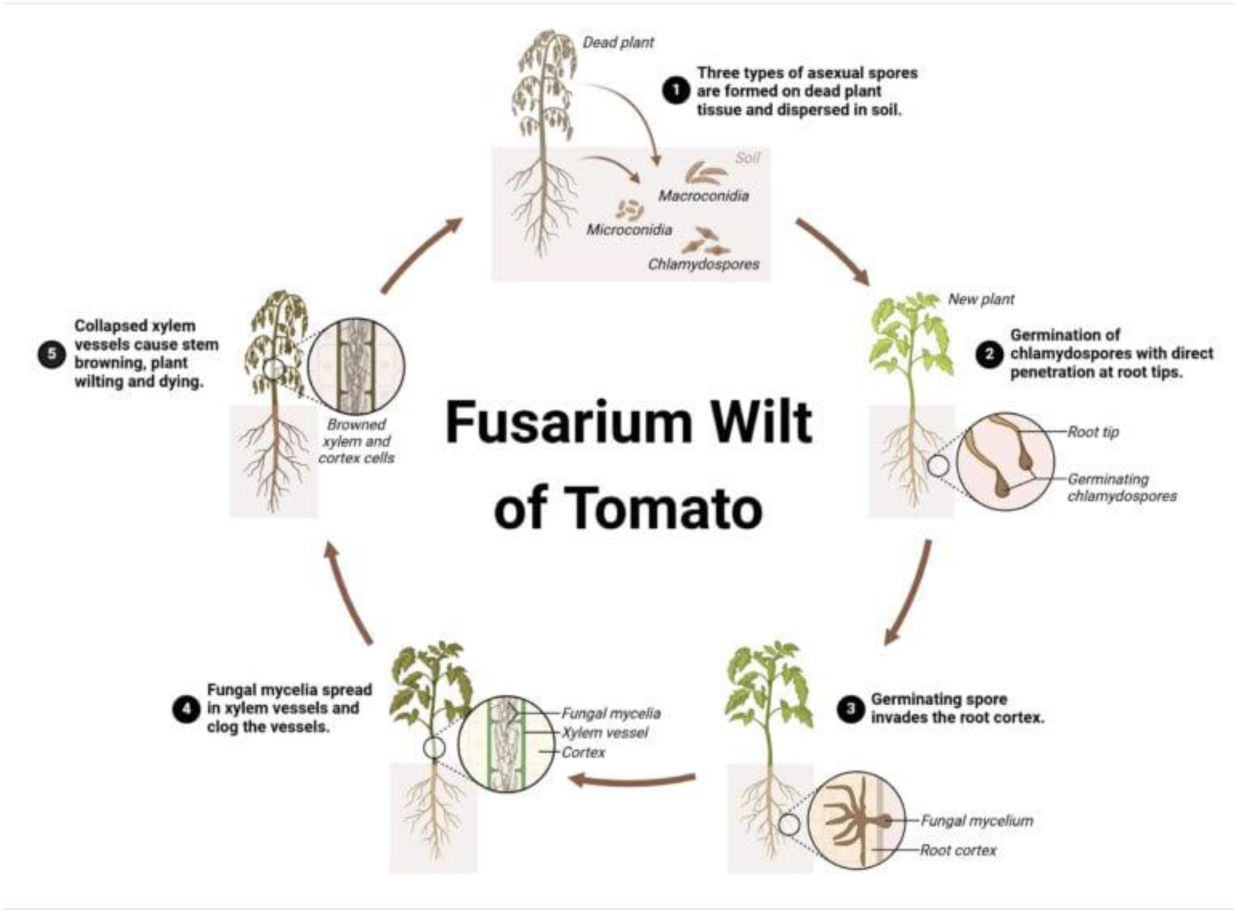
Diagram of the Disease Cycle of Fusarium Wilt.

#### 2.3 INCIDENCE OF FUSARIUM WILT IN NIGERIA

Fusarium wilt, caused by *Fusarium oxysporum f. sp. lycopersici* (Fol), is one of the most destructive and widespread diseases of tomato in Nigeria. The disease has been reported across diverse agro-ecological zones of the country, particularly in northern and central regions where tomato cultivation is concentrated. Its occurrence has been closely linked to the intensive production practices adopted by smallholder farmers, the continuous use of susceptible varieties, and environmental conditions favorable to pathogen survival and proliferation Yusuf, *et al*., (2019).

Field surveys conducted in northern Nigeria have consistently documented high disease incidence, often ranging from 30% to 70%, depending on tomato cultivar, season, and management practices (Olorunjuwon, et al., 2020). For instance, in Kano and Kaduna States, where tomato farming contributes significantly to local livelihoods, Fusarium wilt has been observed as a recurrent problem, particularly in irrigated systems where continuous monocropping allows the pathogen to build up in the soil. Similarly, in Katsina State, the disease incidence was reported to be as high as 60% in fields planted with local susceptible varieties, highlighting the vulnerability of traditional production systems (Yusuf et al., 2019).

In southwestern Nigeria, the incidence of Fusarium wilt is also on the rise, attributed largely to poor soil management, inadequate crop rotation, and reliance on chemical inputs that fail to control soil-borne pathogens effectively (Abayomi, et al., 2021). Smallholder farmers in this region often lack access to resistant tomato cultivars, further increasing the spread and persistence of the disease. This trend demonstrates that the disease is not restricted to northern production zones but is becoming a nationwide challenge.

The persistence of Fusarium wilt in Nigerian tomato fields is largely due to the ability of the pathogen to survive in the soil for extended periods as chlamydospores, even in the absence of a host crop (Agrios, 2005). Such survival capacity allows the disease to remain endemic once established in an area. Additionally, the introduction of contaminated planting material and poor sanitation practices contribute to its wide distribution across different farming communities (Haruna, Yahuza, & Tijjani, 2024).

Overall, the incidence of Fusarium wilt in Nigeria represents a significant threat to sustainable tomato production. Its widespread occurrence across major tomato-producing regions underscores the urgent need for region-specific epidemiological studies and the development of integrated disease management strategies. Without effective intervention, the continued spread of the disease will further undermine food security and the livelihoods of smallholder farmers.

#### 2.4 SEVERITY OF FUSARIUM WILT IN NIGERIA

Fusarium wilt, caused by *Fusarium oxysporum f. sp. lycopersici* (Fol), is not only widespread in Nigeria but also highly severe, resulting in substantial economic and food security challenges. Severity refers to the degree of damage caused by the pathogen on infected plants, often expressed in terms of yield reduction and the intensity of visible symptoms such as wilting, chlorosis, and vascular discoloration. In tomato production systems across Nigeria, severity levels vary according to location, season, and cultivar susceptibility (Haruna, et al., 2024).

Studies in northern Nigeria have shown that severity levels can reach up to 90% yield loss in susceptible tomato cultivars under favorable environmental conditions (Yusuf, et al., 2019). In Kaduna and Kano States, where tomato cultivation is a key agricultural activity, continuous monocropping and reliance on traditional practices exacerbate disease severity (Olorunjuwon, et al., 2020). For smallholder farmers in these regions, severe outbreaks often lead to near-total crop failure, resulting in major financial losses and reduced household food security.

In southwestern Nigeria, similar patterns of severity have been observed, with reports indicating high disease pressure in fields where poor soil fertility, lack of crop rotation, and limited access to resistant varieties prevail (Abayomi, et al., 2021). The severity of Fusarium wilt in these regions has been attributed to both biotic and abiotic factors, including soil type, irrigation practices, and high temperatures, which favor rapid disease development.

The persistence of Fol in soil as chlamydospores further contributes to the long-term severity of the disease. Once established in a field, the pathogen can cause repeated outbreaks for many years, even in the absence of continuous tomato cultivation (Agrios, 2005). This explains why farmers in endemic areas report recurring crop losses despite changes in cropping seasons or partial control efforts. The inability of chemical fungicides to effectively suppress the disease also compounds the problem, leaving farmers with limited management options.

Overall, the severity of Fusarium wilt in Nigeria poses a serious threat to tomato production. Its impact extends beyond yield losses to affect household incomes, food availability, and rural livelihoods. Understanding the severity patterns across different regions is therefore critical for guiding the development of integrated disease management strategies and the introduction of resistant tomato varieties that can withstand the pathogen’s impact.

#### 2.5 ENVIRONMENTAL AND SOCIO-ECONOMIC FACTORS INFLUENCING FUSARIUM WILT IN NIGERIA

Fusarium wilt, caused by the soil-borne pathogen *Fusarium oxysporum,* is a major constraint to the production of key Nigerian crops like tomatoes, bananas, and pulses. Its prevalence and impact across the country are shaped by a confluence of distinct environmental conditions and deeply rooted socio-economic realities faced by local farmers.

##### 2.5.1 Environmental Factors

The Nigerian climate, particularly its high temperatures, creates an ideal environment for the proliferation of *Fusarium oxysporum*. Research has shown that the pathogen thrives in the warm soils common throughout the country, which accelerates disease development and severity (Amusa & Adegbite, 2007). Furthermore, poor soil management practices, including monocropping and a decline in the use of organic amendments, lead to degraded soil health, reducing microbial antagonism and increasing plant susceptibility. Irregular rainfall patterns and drought stress, which are becoming more frequent, also weaken host plants, facilitating fungal invasion and the expression of wilt symptoms (Emechebe & Lagoke, 2002).

##### 2.5.2 Socio-Economic Factors

Socio-economic constraints significantly hinder effective disease management. The high cost and limited availability of certified disease-resistant varieties mean that the majority of smallholder farmers rely on saving and exchanging uncertified seeds or suckers, which is a primary vector for introducing and spreading the pathogen into new fields (Okechukwu & Dixon, 2009). The economic necessity of continuous monocropping of high-value crops like tomatoes to meet market demand leads to a dangerous buildup of inoculum in the soil over time.

Limited access to credit and extension services prevents farmers from investing in integrated pest management (IPM) strategies or rotating with non-host crops, which are often less profitable. This creates a cycle of vulnerability where farmers, aware of the disease, lack the financial capacity or knowledge to implement long-term control measures (Ojiewo et al., 2013). Consequently, Fusarium wilt causes substantial annual yield losses, exacerbating poverty and threatening food security at both household and national levels. Effective management therefore requires strategies that are not only technically sound but also economically feasible and socially acceptable for Nigerian farmers.

#### 2.6 MANAGEMENT APPROACHES TO FUSARIUM WILT

The control of *Fusarium oxysporum f. sp. lycopersici* (Fol), the causal pathogen of Fusarium wilt of tomato, remains a major challenge because of its ability to persist in soil for long periods through chlamydospores and its evolving pathogenic races. A variety of strategies cultural, biological, and genetic have been employed with varying degrees of success.

Cultural practices such as crop rotation, soil sterilization, and organic soil amendments have been reported to lower disease incidence by reducing inoculum levels in the soil (Mia, Shamsi, & Hossain, 2016). Rotation with non-host crops improves soil microbial balance and reduces the pathogen’s survival, though its success is limited by the wide host range of Fusarium species (Indi, 2017). Similarly, organic amendments such as compost and poultry manure have been shown to suppress pathogen activity while enhancing soil fertility (Amponsah, 2015). However, many smallholder farmers in Nigeria rarely adopt these practices due to resource constraints and lack of extension guidance.

Biological control strategies are increasingly recognized as eco-friendly alternatives. Beneficial microorganisms such as *Bacillus subtilis* and *Pseudomonas putida* have demonstrated antagonistic activity against Fusarium through antibiosis, competition, and induction of systemic resistance (Nandini et al., 2017). In addition, *Trichoderma harzianum* has been reported to parasitize Fusarium and improve plant vigor, though the adoption of biocontrol agents remains limited in Nigeria due to poor awareness and lack of commercial formulations (Singh et al., 2018).

Genetic resistance remains the most reliable management option. Resistant tomato cultivars have been developed and used in many parts of the world, offering effective protection against specific races of Fol (Jones, 2016). However, the emergence of new virulent races often overcomes resistance genes, reducing long-term effectiveness. In Nigeria, the availability and adoption of resistant cultivars are still low, mainly due to cost and limited access to improved seeds (Musa & Adeoye, 2020).

#### 2.7 KNOWLEDGE GAP

Despite efforts in the control of *Fusarium oxysporum* f. sp. *lycopersici* (Fol), the causal pathogen of Fusarium wilt of tomato, significant knowledge gaps persist. First, there is limited region-specific epidemiological data on Fusarium wilt in Nigeria, which hampers the development of tailored management practice (Ojo & Fagbola, 2019).

Second, molecular characterization of Fol strains in Nigeria remains underexplored, restricting the design of durable resistance strategies. Third, socio-economic challenges such as inadequate farmer training, weak extension services, and unstable tomato markets discourage the adoption of integrated management approaches (Akinyele, 2021). Addressing these gaps requires multidisciplinary research, stronger policy support, and greater investment in farmer education and resistant seed dissemination.

## CHAPTER THREE

### 3.0 METHODOLOGY

#### 3.1 Study Area

This study will be carried out in Chikun Local Government Area (LGA) of Kaduna State, Nigeria. Chikun lies between latitude 10°20′N and longitude 7°30′E, with an estimated land area of 4,645 km² and a projected population of 368,250 people (NPC, 2006). The area falls within the Northern Guinea Savannah ecological zone, characterized by a tropical climate with distinct wet and dry seasons. The rainy season spans from May to October, with annual rainfall of 1,200–1,600 mm, while the dry season lasts from November to April, with mean daily temperatures ranging from 18°C to 32°C (Kaduna State Bureau of Statistics (KSBS), 2020).

Farming is the primary occupation of the people, with crops such as maize, sorghum, groundnut, and vegetables cultivated extensively. Tomato (*Solanum lycopersicum* L.) is an important cash crop in the area, grown under both rainfed and irrigated conditions, which makes Chikun a suitable location for assessing the incidence and severity of Fusarium wilt (Akanbi et al., 2019).

**Figure 2.**
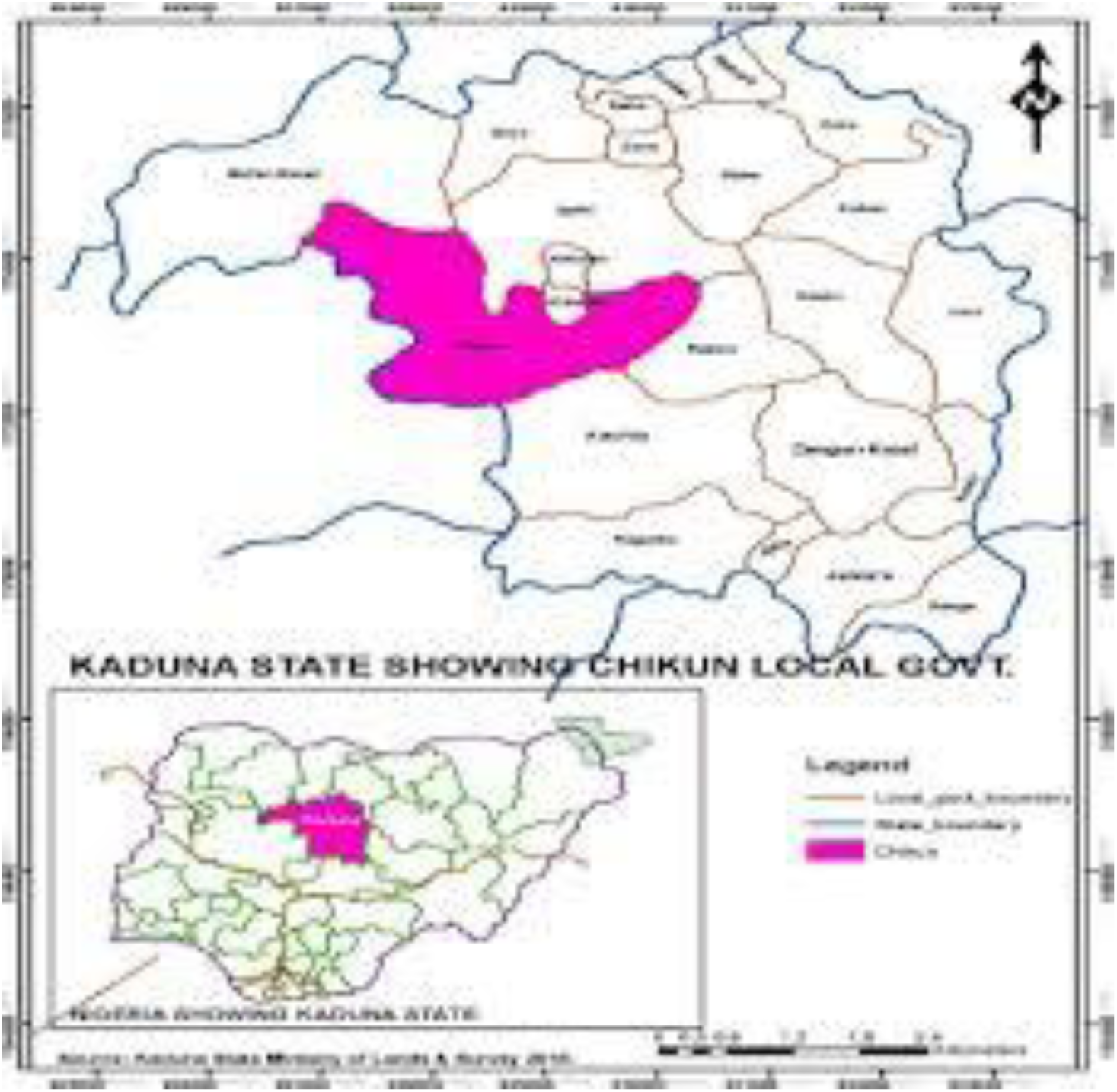
Digital geographical map of Kaduna State showing Chikun local government (SOURCE: Google Earth, 2016)

#### 3.2 Sampling Technique and Sample Size

Sampling is a critical component in research because it allows researchers to gather data from a manageable subset of the population, making the study feasible and efficient (Kumar, 2019). Sampling allows for statistical analysis of data, which is essential for identifying trends and generalizing findings.

A total of three tomato farms (designated as Farms A, B, and C) were surveyed across the selected villages. On each farm, five samples each of tomato stems, leaves, and fruits were collected using a random sampling technique, giving a total of 45 samples (15 per tissue type). Sampling was conducted during the mid to late growing season (July to October), a period when symptoms of Fusarium wilt are most pronounced.

Plants were examined in the field for visual symptoms of Fusarium wilt, including yellowing of lower leaves, one-sided wilting, vascular browning, and stunted growth. Plants were examined for symptoms such as wilting, vascular discoloration, yellowing, and stunted growth, and samples was collected for laboratory confirmation *of Fusarium oxysporum* f. sp. *lycopersici* (Yusuf, Ibrahim, & Adamu, 2019). This random sampling approach ensured representation across al**l** farms studies **a**nd provided a reliable sample size for statistical analysis Yusuf, et al., (2019).

**Figure 3:**
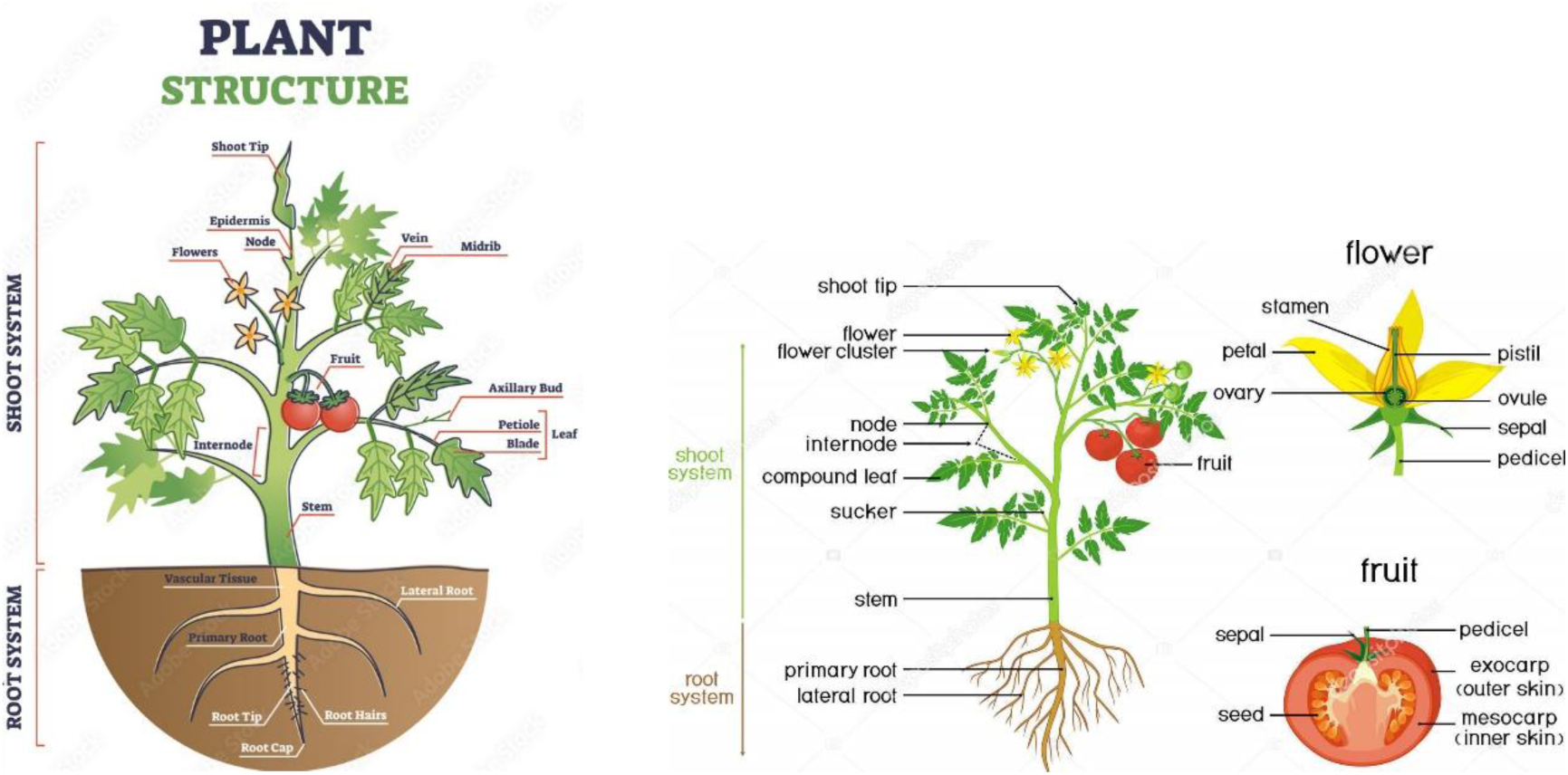
Labeled Diagrams of a Complete Tomato Plant.

#### 3.3 Method of Data Collection

Both primary and secondary data were utilized. Primary data will be collected through;

i. Field surveys and visual inspection of plants for disease symptoms.
ii. Sample collection of tomato stems, leaves, and fruits showing symptoms of wilt.

Secondary data were collected from agricultural extension agents, published research studies, and government agricultural records (Haruna et al., 2024). In the laboratory, we isolated and identified the fungus Fusarium oxysporum f. sp. lycopersici from infected plant tissues using standard methods (Olorunjuwon, et al., 2020). We collected and labeled diseased tomato samples, then analyzed them in the lab to identify the pathogen, following established protocols (Olorunjuwon et al., 2020). The symptoms we observed and recorded were:”

i. One-sided wilting of leaves
ii. Yellowing of lower leaves
iii. Vascular browning visible on longitudinally split stems

Each plant was scored as either infected (1) or healthy (0) based on visual inspection.

##### 3.3.1 Disease Incidence Determination

Disease incidence was calculated using the formula (Okoro et al., 2022):

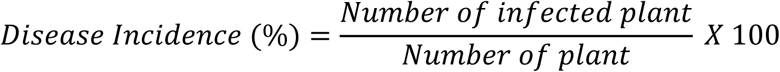

This provided the percentage of tomato plants infected with *Fusarium wilt* across the surveyed farms.

Disease severity Scale was used to assess Fusarium wilt in tomatoes:

| Severity Score | Symptoms Description |
| --- | --- |
| 0 | No symptoms, healthy plant |
| 1 | 1-25% of leaves wilted or yellowed |
| 2 | 26-50% of leaves wilted or yellowed |
| 3 | 51-75% of leaves wilted or yellowed |
| 4 | 76-100% of leaves wilted or yellowed, plant stunted |
| 5 | Plant dead or nearly dead |

Calculating Disease Severity

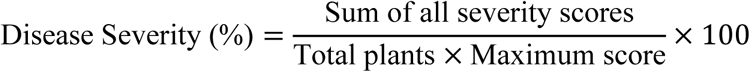

This scale assesses the progression of Fusarium wilt symptoms, from no symptoms (0) to plant death (5).

#### 3.4 Sample Collection and Laboratory Diagnosis

Symptomatic tomato plants were collected from the three surveyed farms. Stems, leaves, and fruits showing typical Fusarium wilt symptoms were excised, labeled according to their respective farms, and placed in sterile paper bags. Samples were then transported to the Plant Pathology Laboratory of the National Veterinary Research Institute (NVRI), Vom, for fungal isolation and identification.

##### 3.4.1 Isolation Procedure

Small sections (5 mm) of the symptomatic tissues were surface-sterilized using 1% sodium hypochlorite (NaOCl) for two minutes, rinsed three times in sterile distilled water, and blotted dry using sterile paper towels. The sterilized sections were plated onto Potato Dextrose Agar (PDA) in sterile Petri dishes. All operations were carried out under aseptic conditions using sterile hand gloves and inoculating tools.

Plates were incubated at 25°C for 5–7 days and examined daily for fungal growth. Colonies with typical *Fusarium* characteristics cottony white to pinkish growth with a violet or purple reverse were sub-cultured onto fresh PDA plates to obtain pure cultures.

##### 3.4.2 Microscopic Identification

Pure cultures were stained with lactophenol cotton blue and examined under a compound microscope. Diagnostic features observed included sickle-shaped macroconidia, oval microconidia, and chlamydospores, consistent with descriptions of *Fusarium oxysporum* f. sp. *lycopersici* by (Leslie and Summerell 2020) and (Elshafie et al., 2021).

The identification confirmed the presence of *Fusarium oxysporum* f. sp. *lycopersici* as the causal organism of Fusarium wilt in the sampled tomato plants. Stems showing characteristic vascular browning yielded the highest isolation frequency, followed by leaves and fruits, which showed moderate to low isolation rates.

#### 3.5 Method of Data Analysis

Data obtained from field and laboratory studies were analyzed using descriptive statistics. The percentage incidence of *Fusarium wilt* was computed for each farm and sample type (stem, leaf, fruit). The results were summarized using tables to show the distribution of infection.

To determine whether differences in infection rates among farms or plant parts were statistically significant, data were subjected to a Chi-square (χ²) test at a 5% significance level (p < 0.05) using the Statistical Package for Social Sciences (SPSS version 25). The analysis helped to evaluate the relationship between farm location and disease incidence

## CHAPTER FOUR

### 4.0 DATA ANALYSIS PRESENTATION AND DISCUSSION OF RESULTS

#### 4.1 Introduction

This chapter presents the results obtained from both field observations and laboratory analyses of tomato samples collected from farms in Kujama, Kakau, and Rido communities of Chikun Local Government Area (LGA), Kaduna State. The findings include laboratory identification of *Fusarium oxysporum* f. sp. *lycopersici*, disease incidence across farm locations and plant parts, and comparative analyses of infection rates. Results are presented in tables, accompanied by interpretations and relevant literature support.

#### 4.2 DATA ANALYSIS PRESENTATION

Table 4.1 presents the frequency of *Fusarium oxysporum* f. sp. *lycopersici* (FOL) isolated from tomato stem, leaf, and fruit samples collected from farms across Chikun LGA. Out of the 45 samples analysed, the pathogen was successfully isolated from 25 samples, representing an overall isolation frequency of 55.6%. Among the plant parts examined, the stem recorded the highest isolation frequency of 66.7%, followed by the leaf with 60.0%, while the fruit had the lowest frequency of 40.0%.

**Table 4.1:** Frequency of Fusarium oxysporum f. sp. lycopersici isolated from tomato samples in Chikun LGA.

| Sample Type | Number Examined | Number of Positive Isolates | Isolation Frequency (%) | Colony/Microscopic Characteristics |
| --- | --- | --- | --- | --- |
| Stem | 15 | 10 | 66.7 | Cottony white to pink colony, purple reverse; macroconidia curved and sickle-shaped; abundant chlamydospores |
| Leaf | 15 | 9 | 60.0 | Whitish colony with sparse aerial mycelia; oval microconidia formed in false heads |
| Fruit | 15 | 6 | 40.0 | Light pink colony, sparse sporulation; few macroconidia and microconidia |
| Total | 45 | 25 | 55.6 | — |

**Source:** Laboratory diagnosis, NVRI Vom (2025).

The high isolation frequency in stem tissues reflects the pathogen’s affinity for vascular tissues, which serve as the main channel for colonization and symptom expression. The results therefore validate field observations of wilting, yellowing, and vascular discoloration as reliable indicators of Fusarium wilt infection. This underscores the importance of regular disease surveillance and pathogen detection in tomato-growing areas to facilitate early control and prevent widespread outbreaks.

Table 4.2 presents the incidence of *Fusarium oxysporum* f. sp. *lycopersici* infection across three tomato farms Kujama, Kakau, and Rido in Chikun Local Government Area (LGA). Out of the 45 tomato plants examined, 27 were infected, representing an overall disease incidence of **60.0%**. Among the farms, Rido recorded the highest disease incidence of 80.0%, followed by Kujama with 60.0%, while Kakau had the lowest incidence of 40.0%.

**Table 4.2:** Incidence of Fusarium wilt by farm location in Chikun LGA.

| Farm Location |  | Number of Plants Examined | Number of Plants Infected | Disease Incidence (%) |
| --- | --- | --- | --- | --- |
| Farm A (Kujama) |  | 15 | 9 | 60.0 |
| Farm B (Kakau) |  | 15 | 6 | 40.0 |
| Farm C (Rido) |  | 15 | 12 | 80.0 |
| **Total/Average** |  | **45** | **27** | **60.0** |

**Source:** Field and laboratory survey, 2025.

The high infection rate observed in Rido may be attributed to predisposing environmental and agronomic factors such as poor drainage, high soil moisture, or continuous tomato cultivation on the same farmland, which favors the survival and proliferation of *Fusarium oxysporum* in the soil. Similarly, the moderate incidence in Kujama and lower level in Kakau could be due to variations in soil fertility, pH, or management practices such as crop rotation and the use of fungicides.

The overall mean incidence of 60.0% indicates that Fusarium wilt is widespread across tomato-producing areas of Chikun LGA. This underscores the need for region-wide management strategies, including crop rotation and the use of resistant varieties, to curb the spread and impact of Fusarium wilt in tomato production systems within Chikun LGA.

Table 4.3 shows that Fusarium wilt affected tomato plant parts at different levels of intensity. Stems recorded the highest incidence (80.0%) and severity (76.0%), with pronounced symptoms such as vascular browning and strong wilting, indicating that the pathogen primarily colonizes the stem’s vascular tissues. Leaves showed moderate incidence (60.0%) and severity (54.0%), with yellowing and partial wilting typical of secondary spread from the stem. Fruits had the lowest incidence (40.0%) and severity (38.0%), displaying only mild discoloration and slight shriveling due to their limited vascular connection. Overall, the results confirm that Fusarium wilt is more severe in tissues with greater vascular exposure, leading to higher symptom expression in stems compared to leaves and fruits.

**Table 4.3:** Incidence of *Fusarium oxysporum* f. sp. *lycopersici* by plant part.

| **Plant Part** | **No. of Samples Examined** | **No. Infected** | **Incidence (%)** | **Mean Severity Score (0–5)** | **Severity (%)** | **Symptom Description (Based on Severity Score)** |
| --- | --- | --- | --- | --- | --- | --- |
| **Stem** | 15 | 12 | 80.0 | 3.8 | 76.0 | Severe vascular browning, longitudinal streaking, intense wilting, partial stem collapse (Severity Class 4: 51–75% wilting). |
| **Leaf** | 15 | 9 | 60.0 | 2.7 | 54.0 | Moderate leaf yellowing, unilateral wilt, marginal necrosis, downward curling (Severity Class 3: 26–50% leaf area affected). |
| **Fruit** | 15 | 6 | 40.0 | 1.9 | 38.0 | Mild discoloration |

| Plant Part | No. of Samples Examined | No. Infected | Incidence (%) | Mean Severity Score (0–5) | Severity (%) | Symptom Description (Based on Severity Score) |
| --- | --- | --- | --- | --- | --- | --- |
|  |  |  |  |  |  | n, slight shriveling, small necrotic patches near pedicel (Severity Class 2: 11–25% area affected). |
| **Total/Mean** | **45** | **27** | **60.0** | **2.8** | **56.0** | — |

**Source:** Laboratory analysis, 2025.

The Chi-square analysis presented in Table 4.4 evaluated whether there were significant differences in Fusarium wilt incidence across farm locations and among different tomato plant parts in Chikun LGA. The results showed that for farm location, the Chi-square (χ²) value was 1.32 with 2 degrees of freedom (df) and a p-value of 0.28, while for plant parts, the χ² value was 2.47 with 2 degrees of freedom (df) and a p-value of 0.18.

**Table 4.4:**
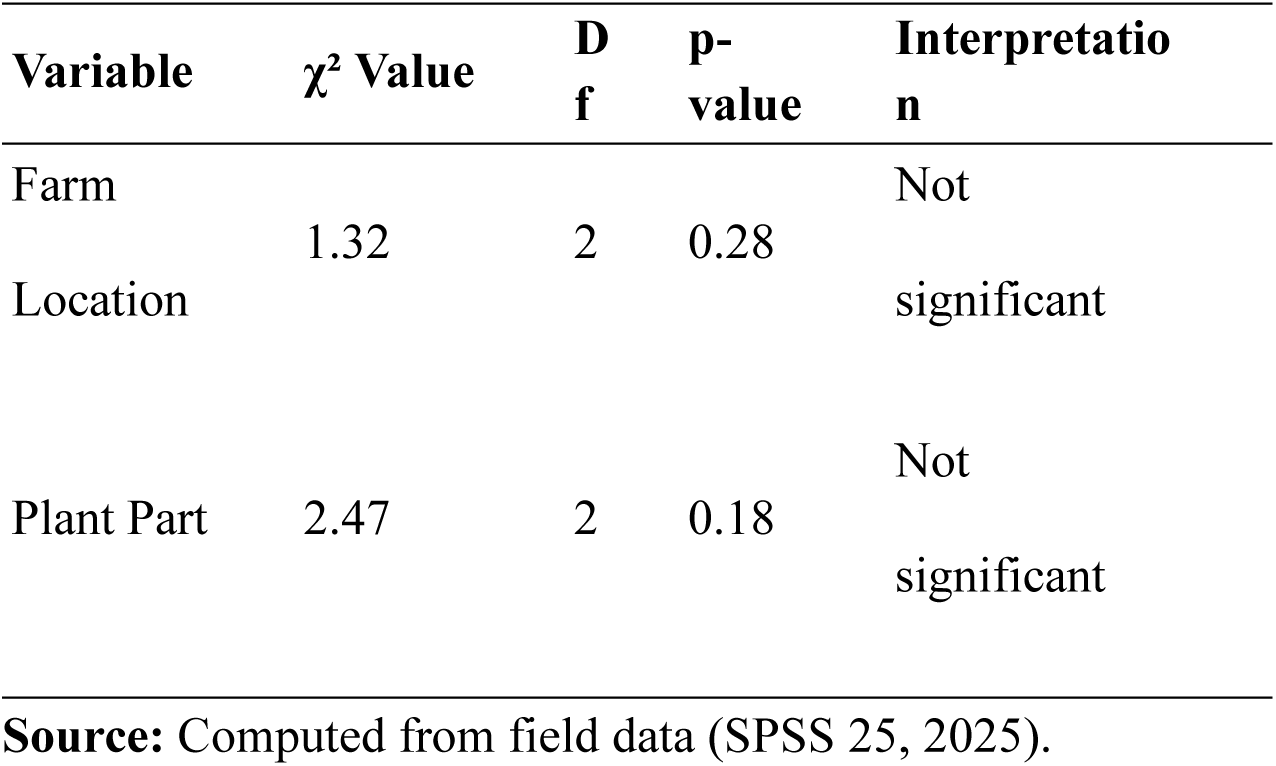
Chi-square analysis of Fusarium wilt incidence among farms and plant parts.

In both cases, the p-values were greater than 0.05, indicating that the observed variations in disease incidence among farms and plant parts were not statistically significant at the 5% level. This implies that Fusarium wilt occurrence was relatively uniform across all sampled farms and tomato plant parts within the study area.

The lack of significant difference suggests that the pathogen *Fusarium oxysporum* f. sp. *lycopersici* is well established and widely distributed in the soils of Chikun LGA, irrespective of farm location or tomato plant organ. This uniform distribution may be attributed to the use of similar agricultural practices, such as continuous tomato cultivation, shared seed sources, and comparable soil conditions, all of which promote the persistence and even spread of the pathogen across fields.

Therefore, the Chi-square analysis confirms that the prevalence of Fusarium wilt in Chikun LGA is widespread and not localized to any specific farm or plant part, emphasizing the need for area-wide integrated management strategies rather than isolated farm-level interventions.

#### 4.2 Laboratory Confirmation of the Pathogen

Microscopic examination of isolates confirmed the presence of *Fusarium oxysporum* f. sp. *lycopersici*. The fungal cultures exhibited cottony white to violet colonies on potato dextrose agar (PDA) and produced diagnostic structures such as sickle-shaped macroconidia, oval microconidia, and thick-walled chlamydospores. These features are consistent with standard descriptions of the pathogen reported by (Leslie and Summerell 2017) and (Elshafie et al., 2021). The consistent recovery of *F. oxysporum* from symptomatic tomato tissues across all farms verified that Fusarium wilt was the major cause of tomato decline in the study area.

#### 4.3 Summary of Results

The study successfully isolated and identified *Fusarium oxysporum* f. sp. *lycopersici* as the causal organism of Fusarium wilt in tomato farms within Chikun LGA. Laboratory analysis of 45 plant tissue samples (stems, leaves, and fruits) revealed typical morphological features of the pathogen, including white to pinkish cottony colonies and sickle-shaped macroconidia. The overall disease incidence recorded across the three surveyed farms was 60.0%, with Rido showing the highest infection rate (80.0%), followed by Kujama (60.0%) and Kakau (40.0%). Among plant parts, the stem had the highest incidence (80.0%), followed by the leaves (60.0%) and fruits (40.0%). Statistical analysis indicated no significant difference (p > 0.05) in infection rates among farms and plant parts, confirming the widespread nature of the disease.

#### 4.4 Discussion of findings

The present study confirmed the occurrence of *Fusarium oxysporum* f. sp. *lycopersici* (FOL) as the causative agent of tomato wilt in three communities of Chikun LGA. The laboratory analysis revealed an overall isolation frequency of 55.6%, indicating that more than half of the tomato samples collected were infected by the pathogen. Among the plant parts examined, the stem recorded the highest isolation frequency (66.7%), followed by leaves (60.0%) and fruits (40.0%). This agrees with (Akinbode and Ikotun 2019), who reported that *F. oxysporum* primarily colonizes the vascular tissues of tomato stems, causing wilting due to disruption of water transport. Similarly, (Nirmaladevi et al., 2020) observed that the pathogen thrives within xylem vessels, resulting in typical wilt symptoms.

The incidence value of 60.0% recorded in this study is consistent with the findings of (Musa et al., 2021), who reported a 62% incidence in tomato farms in Adamawa State, and (Bello et al., 2020), who recorded 58% in Gombe State. Likewise, (Abdullahi et al., 2019) reported 50% incidence among smallholder farmers in Kano, emphasizing that Fusarium wilt remains a persistent threat to tomato production in northern Nigeria. The relatively high incidence in Chikun may be due to continuous tomato cultivation on the same farmland, poor crop rotation, and the use of contaminated seeds, as also noted by (Okungbowa and Ebhodaghe 2018).

The non-significant chi-square results suggest that the disease is uniformly distributed across farms and plant parts. This aligns with (Osei et al., 2020), who observed similar uniform distribution in Ghana, attributing it to the widespread presence of contaminated soil and seedborne inoculum. Compared to the 68% incidence reported by (Ibrahim et al., 2022) in Plateau State, the moderate incidence in Chikun may be influenced by microclimatic and soil conditions, as (Aguoru et al., 2019) highlighted that warm and humid environments favor pathogen proliferation.

Morphological identification confirmed characteristic *F. oxysporum* features such as macroconidia, microconidia, and chlamydospores, consistent with (Leslie and Summerell 2017). Although morphological identification is reliable, (Nelson et al., 2018) and (Elshafie et al., 2021) recommend molecular techniques for precise race identification. Future studies should therefore include polymerase chain reaction (PCR) assays to determine pathogenic races and support breeding for resistance.

The persistence of the disease in Chikun likely results from limited farmer awareness and inadequate field sanitation, as observed by (Ezekiel and Oladiran 2018). The findings thus emphasize the need for integrated disease management (IDM) approaches, including crop rotation, use of resistant varieties, and biological control. (Babalola et al., 2021) reported that crop rotation with cereals such as maize effectively reduces soil inoculum levels, while (Sani et al., 2020) demonstrated that combining resistant cultivars with organic amendments suppresses Fusarium wilt.

Overall, the results confirm that *Fusarium oxysporum* f. sp. *lycopersici* is a major pathogen limiting tomato production in Chikun LGA. Similar to previous studies in northern Nigeria, the disease’s widespread occurrence calls for urgent management strategies integrating resistant varieties, good field hygiene, and farmer training to ensure sustainable tomato production.

## CHAPTER FIVE

### 5.0 SUMMARY OF FINDINGS, CONCLUSION AND RECOMMENDATION

#### 5.1 Summary of Findings

The study investigated the prevalence, incidence, and distribution of Fusarium wilt (*Fusarium oxysporum* f. sp. *lycopersici*) affecting tomato (*Solanum lycopersicum* L.) production in Chikun Local Government Area (LGA) of Kaduna State, Nigeria. Findings revealed that *F. oxysporum* was successfully isolated from 55.6% of the total tomato samples analyzed, confirming its presence as the primary causal agent of tomato wilt in the study area. The overall disease incidence was recorded at 60.0%, signifying a moderate-to-high level of infection across the surveyed farms. Among the three farms studied, Rido exhibited the highest incidence (80.0%), followed by Kujama (60.0%) and Kakau (40.0%).

Analysis by plant part showed that the stem had the highest infection frequency (80.0%), followed by leaves (60.0%) and fruits (40.0%), reflecting the vascular nature of the pathogen and its tendency to colonize xylem tissues. Statistical evaluation using chi-square analysis indicated no significant differences (p > 0.05) in disease incidence among farms and plant parts, suggesting uniform distribution of the pathogen across the study area. Morphological characterization of isolates revealed cottony white to pink colonies with purple reverses, sickle-shaped macroconidia, and chlamydospores, which are typical diagnostic features of *F. oxysporum* f. sp. *lycopersici*. The study further revealed that most farmers practiced continuous tomato cultivation on small plots, often using susceptible varieties and minimal disease management techniques. These findings collectively demonstrate that Fusarium wilt is widespread in tomato-producing areas of Chikun LGA and represents a significant constraint to tomato yield and productivity.

#### 5.2 Conclusion

The findings of this study conclusively establish that *Fusarium oxysporum* f. sp. *lycopersici* is a major pathogen responsible for tomato wilt in Chikun Local Government Area of Kaduna State. The consistent isolation of the fungus from infected plant tissues, coupled with its distinctive morphological characteristics, confirmed its role as the causal agent of the disease. The overall incidence of 60.0% recorded across the surveyed farms indicates that Fusarium wilt is both prevalent and persistent within the area’s tomato production systems.

The high infection rate observed in stem tissues compared to leaves and fruits reflects the vascular colonization habit of the pathogen, which disrupts water and nutrient translocation, leading to typical wilting symptoms. The non-significant variation in disease incidence among the different farm locations suggests that the pathogen is uniformly distributed across tomato-growing areas in Chikun LGA, possibly due to continuous cultivation, contaminated soils, and the use of infected planting materials.

This study demonstrates that inadequate crop rotation, poor field sanitation, and limited awareness among farmers have contributed to the persistence and spread of Fusarium wilt in the study area. Therefore, urgent and coordinated management interventions are required to reduce yield losses and sustain tomato production. Such measures should include the adoption of resistant cultivars, implementation of proper crop rotation schemes, biological control, and farmer education on effective disease management practices.

#### 5.3 Recommendations

Based on the findings and conclusions of this study, the following recommendations are proposed to reduce the prevalence and impact of *Fusarium oxysporum* f. sp. *lycopersici* on tomato production in Chikun LGA:

1. Farmers should cultivate tomato varieties that are genetically resistant or tolerant to Fusarium wilt. The use of improved resistant cultivars will significantly minimize disease incidence and yield losses.
2. Continuous tomato cultivation on the same farmland should be discouraged. Rotating tomato with non-host crops such as maize, sorghum, or legumes will help break the disease cycle and reduce soil inoculum buildup.
3. Regular farmer education programs should be organized by agricultural extension agents to raise awareness on disease identification, proper sanitation, and the importance of using disease-free planting materials.
4. Farmers should regularly monitor and manage soil pH and moisture, as acidic and poorly drained soils favor pathogen survival and infection. The application of organic matter and lime can help improve soil structure and suppress fungal growth.
5. Government agencies and research institutions should support integrated disease management research, encourage seed certification programs, and promote access to resistant varieties and affordable biocontrol inputs.

#### 5.4 Limitations of the Study

Although this study provided valuable insights into the prevalence and distribution of *Fusarium oxysporum* f. sp. *lycopersici* in Chikun LGA, several limitations were encountered during the research. Firstly, the sample size of forty-five tomato plant samples, though representative, may not fully capture the diversity of the pathogen population or the varying field conditions across the entire local government area. A larger sample size could have provided more comprehensive data for statistical analysis and spatial comparison. Secondly, environmental factors such as soil temperature, relative humidity, and nutrient status were not monitored throughout the cropping season due to limited laboratory resources and time constraints. These factors could have provided a deeper understanding of the ecological drivers influencing disease prevalence and severity. Despite these limitations, the study successfully established a strong baseline for understanding the epidemiology of Fusarium wilt in the study area and offers valuable guidance for further research and management planning.

#### 5.5 Contributions to Knowledge

This study has made several important contributions to existing knowledge on the epidemiology and management of *Fusarium oxysporum* f. sp. *lycopersici* affecting tomato production in northern Nigeria:

i. The study provides updated baseline information on the incidence and distribution of Fusarium wilt in tomato fields across Chikun LGA, thereby filling a major data gap in Kaduna State.
ii. The research established that tomato stems are the most susceptible plant part to Fusarium wilt, confirming the vascular nature of the pathogen’s colonization pattern.
iii. Statistical analysis revealed that Fusarium wilt incidence did not vary significantly among farms and plant parts, demonstrating the widespread and uniform distribution of the disease within the study area.
iv. The study highlighted the role of continuous cropping, poor field sanitation, and limited farmer awareness as contributing factors to the persistence of *F. oxysporum* in tomato-growing soils.
v. The findings provide a scientific foundation for the development of integrated management strategies, including resistant variety selection, biological control, and soil management interventions tailored to local farming systems.

Through these contributions, the study enhances scientific understanding of Fusarium wilt dynamics in smallholder tomato production systems and serves as a reference point for future research, extension activities, and policy formulation on tomato disease management in Nigeria.

#### 5.6 Suggestions for Further Studies

In view of the findings and limitations encountered in this study, the following suggestions are proposed for future research on Fusarium wilt of tomato in Nigeria:

i. Extended research covering multiple cropping seasons should be conducted to evaluate how soil temperature, pH, moisture, and nutrient composition influence the epidemiology of Fusarium wilt under field conditions.
ii. Future research should also examine the economic losses and livelihood implications associated with Fusarium wilt outbreaks to guide policy interventions and prioritize support for smallholder tomato farmers.
iii. Studies should focus on creating simple, affordable diagnostic kits for early detection of *F. oxysporum* in the field, enabling timely disease management and reducing crop losses.

